# Conflict of Interest Policies at German medical schools - A long way to go

**DOI:** 10.1101/809723

**Authors:** Peter Grabitz, Zoe Friedmann, Sophie Gepp, Leonard U. Hess, Lisa Specht, Maja Struck, Sophie Tragert, Tobias Walther, David Klemperer

**Author notes:** Authors contributed equally (Peter Grabitz, Zoe Friedmann, Sophie Gepp, Leonard U. Hess, Lisa Specht, Maja Struck, Sophie Tragert, Tobias Walther).

## Abstract

**Background:** Most medical students are in contact with the pharmaceutical or medical device industry during their studies. Medical schools play an important role in protecting students from undue commercial influence and educating them about pharmaceutical marketing practices. Such influence has been shown to affect later prescribing behaviour with potential adverse effects for patient care. While in North America, many medical schools formulated and implemented conflicts of interest (COI) policies, only few such institutional policies have been reported in Germany. We aimed to analyze the quantity and quality of policies and curricula on COI at medical schools across Germany.

**Methods:** We collected relevant COI policies and teaching activities by conducting a search of the websites of all 38 German medical schools using standardized keywords for COI policies and teaching. Further, we surveyed all medical schools’ dean’s offices and adapted a scoring system for obtained results with 13 categories based on prior similar studies.

**Results:** We identified relevant policies for one medical school via the web-search. The response rate of the deans’ survey was 16 of 38 (42.1%). In total, we identified COI-related policies for 2 of 38 (5.3%) German medical schools, yet no policy was sufficient to address all COI-related categories that were assessed in this study. The maximum score achieved was 12 of 26. 36 (94.7%) schools scored 0. No medical school reported curricular teaching on COI.

**Conclusion:** Our results indicate a low level of action by medical schools to protect students from undue commercial influence. No participating dean was aware of any curriculum or instruction on COI at their respective school. The German Medical Students Association and international counterparts have called for a stronger focus on COI in the classroom. We conclude that for German medical schools there is still a long way to go.

## 1. Introduction

### 1.1 Background

Contacts between pharmaceutical or medical device industry and healthcare professionals have long been a point of discussion, as they may lead to conflicts of interest (COI). According to the widely accepted definition from the Institute of Medicine, COI are regarded as circumstances that create a risk that professional judgments or actions regarding a primary interest will be unduly influenced by a secondary interest (1). In healthcare, COI may exist between the physician’s commitment to patient care and industry’s primary aim to sell their products. This ethical tension presents challenges towards medical professionalism (2) and becomes exacerbated when trust is additionally eroded by failure to disclose commercial ties. For example, recently the prominent breast cancer researcher José Baselga had stepped down from his role as chief medical officer of Memorial Sloan Kettering Cancer Center after failing to report his financial conflicts of interests in multiple research articles (3). However, universities and medical schools are increasingly expected to conduct translational research from “bench to bedside” - a paradigm that includes market commercialization and requires industry collaborations which makes contact with the private sector inevitable. In order to protect independent patient care, professional handling of conflicts of interest by physicians is essential. Therefore, from the very beginning the education of the next generation of physicians should be based on sound clinical evidence, free from bias arising from the commercial interests of industry.

In this context, it is crucial to understand medical school as a pivotal phase for future doctors, which not only provides knowledge through lectures and courses, but also attitudes through the environment it creates. It has been argued, that physicians’ attitudes towards the pharmaceutical industry and their inclination to be influenced by marketing efforts manifest early during their professional training (4). A large body of evidence exists showing that medical students themselves are in contact with industrial companies on a regular basis (4–12). Contacts increase in the course of studies, with more interactions during the clinical part of their studies (5,13,14). A study by Lieb et al. (8) at eight German medical schools revealed that only 12% of surveyed students had never received a gift or attended a sponsored event. The authors also found that 60% of these students had a promotional gift handed on to them by a physician they worked with, who received the gift by a company beforehand (8). Professors and other physicians act as role models students base their attitudes and actions on - not only regarding their clinical work, but also regarding interactions with industry and COI. The actions of those role models constitute a “hidden curriculum” and conceptualize what is perceived to be normal (15). Therefore, the environment shaped by medical schools teaches unsaid lessons on how to deal with industrial marketing.

In academic research, it is common practice to disclose COI as part of the publication process in peer reviewed journals and it has also become a norm to include a separate COI slide during presentations. Nonetheless, financial relationships of teaching staff and faculty with industry are not commonly disclosed to medical students, although these relationships may affect academic and publishing interests, the content they choose to disseminate to medical students and their general professional medical opinions (16,17). The extent to which teaching faculty in Germany has financial ties to industry actors remains largely unclear. Despite frequent debate, there is currently no German equivalent to the Physician Payments Sunshine Act in the United States of America, where information on payments from industry to physicians is collected, categorized and made publicly available by law (18). Data reported by CORRECTIV (19) based on voluntary disclosures indicate that physicians, pharmacists and other healthcare professionals together with their respective institutions received a minimum of 562 million euros in 2016 alone. How many of these providers had teaching responsibilities at medical schools is largely unknown.

### 1.2 The role of conflict of interest policies at medical schools

If contact with the industry is unavoidable for medical students, what actions are currently taken by medical schools to protect students from undue influence and to guarantee unbiased teaching content? Some studies found that 65% of surveyed medical students in Germany (20) and 85.2% in France (9) reported feeling inadequately prepared for interactions with the pharmaceutical industry. 90% of those students in Germany reported that dealing with industrial marketing practices had never been addressed during their lectures (20). Indeed, it remains unclear to what extent German medical schools include COI topics in their curricula. In a survey (21), 14.4% of German medical students who took part in the study noted they took part in a lecture or course dealing with COI; of those classes, however, 90% were optional.

Aside from teaching about industry practices of marketing and promotion, restrictive COI policies at the medical school level have been suggested to increase students’ awareness of the consequences of inappropriate marketing practices in the learning environment (22). Some studies indicate that COI policies at medical schools have a significant impact on prescribing practices and inoculate physicians against persuasive aspects of pharmaceutical promotion (23–25). These studies also show that more restrictive policies were associated with higher compliance of physicians’ prescriptions with current guidelines (23–25). It has not been clarified which components of a COI policy might have a particularly important impact on physicians’ behaviour and attitudes. Prior work assessing the quantity and quality of medical school COI policies was conducted in Australia (26), Canada (27), France (28) and the US (29). Some of this research encouraged change among medical schools so that in 2014, 136 of 160 US medical schools had an existing policy on COI (29). In November of 2017, after a study was published which found no existing policies at any of the French medical schools (28), the French Deans’ Conference of Medicine and Odontology Schools published an “Ethical and deontological Charter” (30) - effectively a COI policy for all French medical schools on a national level. In Germany, Lieb et al. found that in 2013 only two medical schools reported having a COI policy (21). However, neither of both schools reporting a policy (TU Dresden and RWTH Aachen) supplied the policies themselves and hence, the content and strength of the policies remain unclear.

The objective of this study was to determine whether medical schools in Germany have institutional COI policies in place and to rate the strength of the policies obtained by means of 13 pre-defined criteria. As part of these criteria, whether or not medical schools offer curricular teaching activities addressing COI in medicine was also examined.

## 2. Methods

Our methodology built upon criteria used in earlier studies on COI policies such as the American Medical Students Association (AMSA) scorecard (29), the Canadian scorecard by Shnier et al. (27), and the French conflict of interest ranking by Scheffer et al. (28) A list of the 38 German medical schools was obtained from the website of the German Medical Faculty Association (Medizinischer Fakultätentag) (31). After formal exchange with a member of the German Ethics Council about the nature of this study, which only involves policies at an institutional level rather than patient data or other personal information, it was deemed unnecessary to ask for formal approval from an ethics committee.

### 2.1 Web-based search

Two researchers (LS, MS) independently searched the websites of the respective medical schools (or if nonexistent, the websites of the respective universities) using integrated search engines in June 2018 to identify policies related to COI, documents interpreting policies or material published regarding COI in the curriculum. Addresses of the websites searched are listed in the supplementary material. Search terms included “Interessenkonflikt”/“Interessenskonflikt” (conflict of interest), “Industrie” (industry), and “interne Regulierung” (internal regulation) based on previous publications (28). If a policy was in place, it was recorded together with the latest date of review. Only policies that specified their oversight over medical schools were considered relevant for this study. Therefore, policies applying to an entire university or only to a university medical center were excluded. Disagreement about inclusion of the recorded sources was discussed with all authors. Those sources included were later assessed via the methodology previously determined through the scoring criteria in our codebook (as described in ‘results’) (32).

### 2.2 Contacting medical schools

In May 2018, we contacted each office of the dean of medicine to inform them about the study through a written letter. The letter gave background information about the study’s purpose and outlined the criteria for which we needed documentation. We asked the medical school to send any form of policy (or parts of a policy) relating to the management of conflicts of interest, as well as information on enforcement of the policy. Furthermore, the letter íncluded the request to provide information on curriculum contents addressing the consequences and management of COI. We did a maximum of three follow-ups for non-responders. We first sent an email in June 2018 reiterating the content of the letter previously sent. We then followed up via email in July 2018 and enclosed two letters of support, one from David Klemperer and one from Barbara Mintzes, co-author of the study which analyzed conflict of interest policies at Canadian medical schools and editor of a widely-used teaching manual on pharma promotion (33). In August 2018, we followed up by sending the results of the web-based search. Representatives of the dean’s offices were given the opportunity to confirm, correct or comment on our web-based findings. In addition to searching the websites and contacting the offices of the deans of medical schools, we sought information via personal contacts and experts in the field. Data cut-off was October 2018. We excluded policies from affiliated teaching hospitals, because they are not under the authority of the dean of the medical school. Further, we excluded any policies or parts of policies that did not specifically apply to a medical school.

### 2.3 Scoring system

We adapted a scoring system based on criteria used in earlier studies by Scheffer et al. (28) and Shnier et al. (27) in the French and Canadian context respectively, as well as the AMSA Scorecard (29). The following categories were addressed:

1. Gifts from industry
2. Meals from industry
3. Consulting relationships
4. Industry-funded speaking relationships
5. Educational activities like CME-lectures
6. Participation in promotional events
7. Honoraria and scholarships from pharmaceutical industry
8. Ghostwriting and honorary authorships
9. Industry Sales Representatives
10. Disclosure
11. Medical school curriculum on COI
12. Extension of policy
13. Enforcement of policy

Subsequently, we graded the results for each category through our scoring system from 0 to Generally, “0” means no policy or a permissive policy, “1” a moderate policy and “2” a restrictive policy. A German codebook outlining the decision pathway for each category is available online (32). Three reviewers independently (LH, TW, ST) undertook the scoring of the medical schools’ policies. All authors then reviewed the scoring. Any disagreement was resolved through discussion and majority vote. We then summed up the scores of all individual categories for each medical school to create a global score, with a range of 0 to 26 points. No weighting of single categories was performed.

## 3. Results

### 3.1 Web-based search

The web-based search was conducted to identify publicly available COI policies at German medical schools. The search yielded relevant results for one of the 38 medical schools: an anti-corruption brochure and a third-party funds statute from Charité-Universitätsmedizin Berlin (**Fig.1**). Additional articles and publications were identified but excluded from analysis, because they either did not relate to predefined criteria or did not specifically apply to the entire medical school. Our web-based search strategy revealed no information on relevant compulsory curricular teaching activities addressing COI. One elective course at Friedrich-Schiller-Universität Jena was identified.

**Figure 1:**
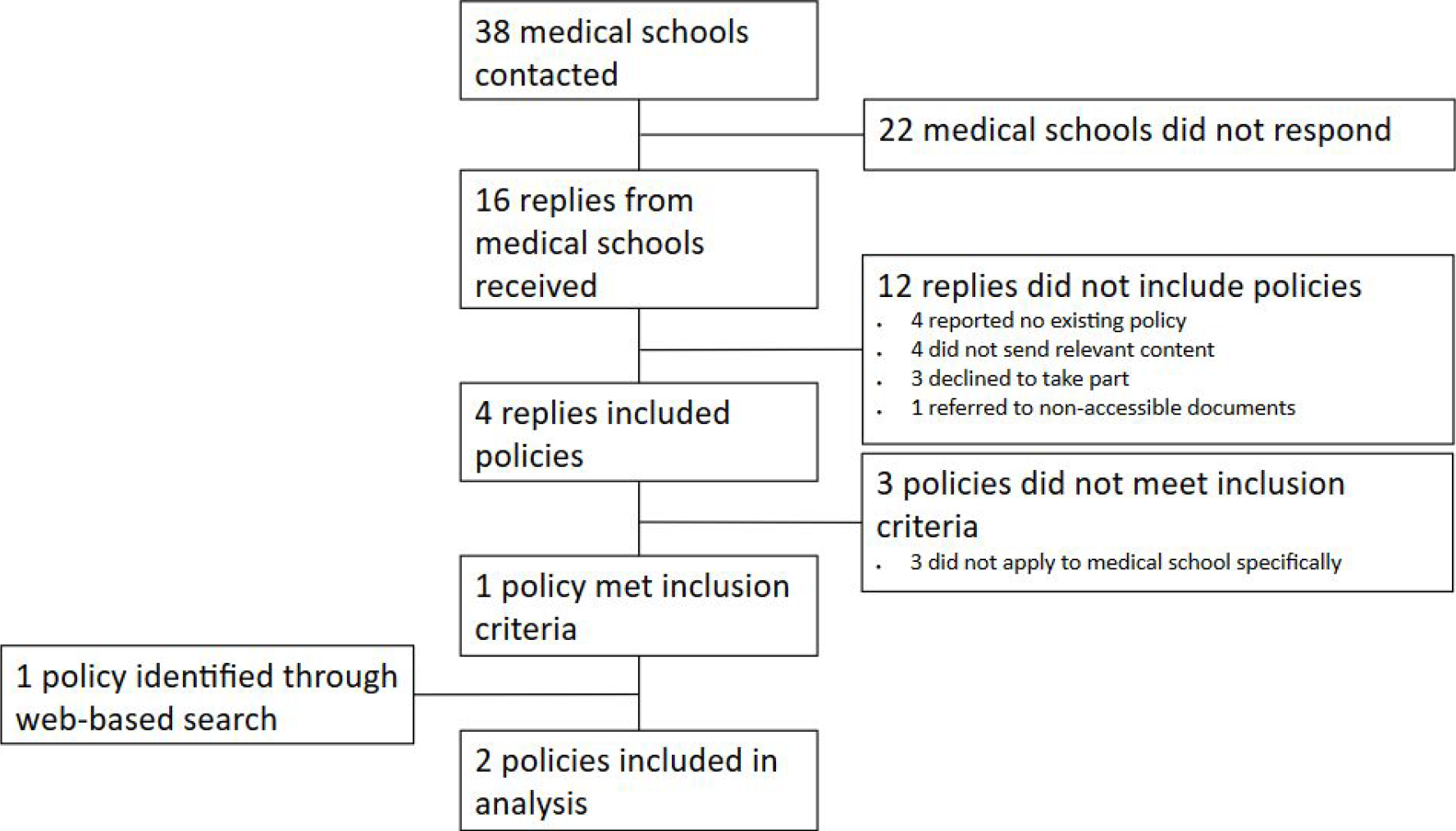
Flowchart of included COI policies

### 3.2 Contacting medical schools

German medical schools were contacted to provide validated insight into existing COI policies. The total response rate was 42.1% (16 of 38). Twelve of the responding medical schools did not send policies. Four medical schools (10.5%) included policies dealing with COI, of which three (an anti-corruption directive and a monetary benefit acceptance policy from the Ludwig-Maximilian-Universität München, a code of practice as well as an anti-corruption directive from the Julius-Maximilians-Universität Würzburg, a compliance brochure, gifts and benefits acceptance policy, and a third-party funds statute from the Friedrich-Schiller-Universität Jena) exclusively applied to university medical centers, not to the respective medical schools, and were therefore excluded from further analysis. One policy met inclusion criteria and comprised an anti-corruption directive issued by the medical school and university medical center of the Technische Universität Dresden (**Fig. 1**).

Of the 16 replies, 5 medical schools (13.2%) (Universität des Saarlandes, Albert-Ludwigs-Universität Freiburg, Georg-August-Universität Göttingen, Christian-Albrechts-Universität zu Kiel, Universität Witten/Herdecke) responded not having COI policies or that COI was not part of the curriculum. The Universität des Saarlandes stated that there was no separate policy for the medical school, while the Albert-Ludwigs-Universität Freiburg declared not having a COI policy within the medical curriculum, as well as no explicit lectures on COI. Also the Christian-Albrechts-Universität zu Kiel reported no existing COI policy within their medical school, neither was the topic taught in the medical curriculum. The reply from the Georg-August-Universität Göttingen stated that basic knowledge about pharmacoeconomics was taught, however, not mentioning corruption and transparency within the medical system. As stated by the Universität Witten/Herdecke, COI management lies with the contracted teaching hospitals. The Friedrich-Alexander-Universität Erlangen-Nürnberg replied that several policies apply within their university, however, no COI policy relevant to this study, issued by the medical school itself is externally available. The Universität Greifswald and the Medizinische Fakultät der Universität Hamburg initially asked for more time to reply, yet did not send material until the end of the data collection period. The Universität Augsburg is still in the process of setting up a medical curriculum, welcoming medical students starting in 2019 and was hence not able to report on COI policies or teaching activities. No further response as to whether a general COI policy existed was received. The Westfälische-Wilhelms-Universität Münster and the Goethe-Universität Frankfurt reported no capacities to take part in our study, while the Justus-Liebig-Universität Gießen actively decided against participating. The Universität Ulm neither addressed COI policies nor curriculum contents in their reply. The remaining medical schools did not respond to any request during the data acquisition period.

### 3.3 Analysis of COI policies

The two included policies were assessed according to a predefined scoring system as set out in our codebook (32). Results of each analysis are listed in **Table 1**. The Technische Universität Dresden achieved the highest score among the medical schools receiving a score of 12 out of 26. Charité Universitätsmedizin Berlin scored 4 points in total. All other medical schools did not supply a valid COI policy and had no retrievable information on COI policies on their websites according to inclusion criteria (**Fig. 1**).

**Table 1:**
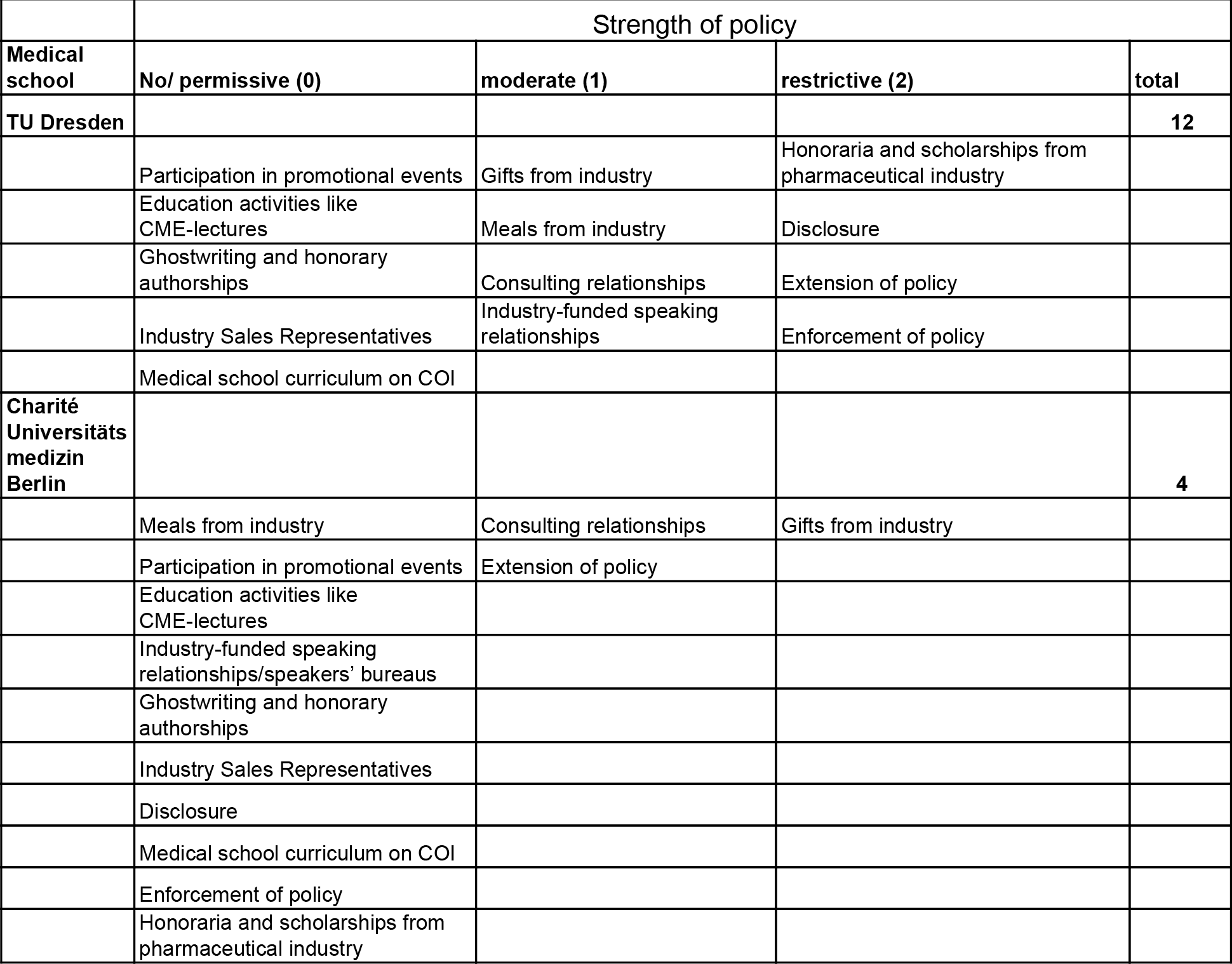
Overview about scoring of policies

We did not acquire any information about obligatory teaching activities on COI through web-based search or the deans survey. However, through personal contacts and seeking advice from experts, we received information on courses that cover COI at 3 medical schools (7.9%): Charité Universitätsmedizin Berlin, Universität Mainz and Universität Leipzig. These teaching activities are either lectures in which COI is discussed (Universität Leipzig, Universität Mainz, Charité Universitätsmedizin Berlin) or elective courses that students can choose within their curriculum (Charité Universitätsmedizin Berlin, Universität Mainz).

## 4. Discussion

Education and policies on COI have been suggested to sensitise medical students against undue influence by industry (5,23–25). Medical students themselves increasingly demand stronger COI regulations, disclosure of teaching faculty’s COI and courses on COI. The German Medical Students’ Association (bvmd e.V.) adopted a position paper on the independence of education in 2013 (34), in May 2019 the European Medical Students Association (EMSA) passed a policy titled “Conflicts of Interest in Medical Education Settings” (35) and the International Medical Students’ Association (IFMSA) followed in August 2019 with a policy called “Integrity and transparency in medical education” (36). These actions are indicative of broader student interest in policy change. Within this study we identified two German medical schools with policies concerning COI. However, none of these policies sufficiently covered the broad spectrum of evaluated categories with relevance to COI, nor did they focus explicitly on medical education. These results indicate little effort by German medical schools to address the issue of COI in medical education.

Only 16 out of 38 German medical schools chose to participate in our study. The best performing policy was an anti-corruption directive issued by the Technische Universität Dresden that included four restrictive and four moderate elements related to different scoring categories. Yet we were unable to retrieve this policy from the medical school’s website during the performed web-search. Prior studies excluded non-public policies from analysis (26,27), since an inaccessible, not widely circulated policy is unlikely to have a relevant impact and may also go unrecognized by academic staff. In this context, a study from 2014 reported that 87.8% of medical students in Germany did not know whether a policy regarding conflicts of interest existed at their school (21). Within this study two medical schools were reported to have a policy on COI (21). Our research could only verify one of those COI policy equivalents at TU Dresden. RWTH Aachen reported a policy in 2014 but did not reply to our study, nor was a policy identified on the school’s website. Despite six medical schools committed to the development of a COI policy (21), our results indicate that no policy has been published since 2014. We furthermore did not receive any information about teaching on COI through the deans survey. This is in contrast to the survey by Lieb et al (21). In their study, deans from seven medical schools reported COI teaching activities (Universität Bonn, Universität Erlangen-Nürnberg, Universität des Saarlandes (Homburg), Universität Gießen, Universität Göttingen, Universität Frankfurt, Universität Köln). From these medical schools, only the dean’s office of Universität Göttingen stated on COI teaching following our request and declared that their curriculum included basic education on pharmacoeconomics but did not explicitly cover COI related aspects like transparency or corruption. We found COI teaching activity at German medical schools, if existent, to be an initiative by singular faculty members rather than a structured component of the curriculum. From personal contacts we learned about courses at Charité Universitätsmedizin Berlin, Universität Mainz and Universität Leipzig, our web-search found one course at Friedrich-Schiller-Universität Jena. These tend to be either singular lectures only or non-mandatory electives, which are not transversally integrated into the curriculum and thus are likely to have limited impact.

Comparable studies to ours were conducted in the United States, Canada, Australia and France (26–29), allowing for an international comparison of our results. In general, North American medical schools tackle the issue of COI in medical education more proactively. In Canada 16 of 17 medical schools had some form of COI policy in place in 2013 (27) and in 2014, 136/160 US medical schools reported an existing policy on COI (29). The Australian study found that 7 medical schools out of 20 had a COI policy. The French study exposed similar results as our own data. They found no formal COI policy at any of the 37 French medical schools and only scattered COI teaching activities. Their response rate of 8,1% may be indicative of the low interest in the topic by medical schools in at this time. The publication of these results led to increased media attention (37) and ultimately the French deans’ conference adopted a nation-wide COI policy (30).

Our results suggest that COI are a neglected topic at medical schools in Germany, particularly with regard to the education of medical students. The scarce efforts to include COI in teaching are all the more surprising, since the German National Competence-Based Learning Objectives for Undergraduate Medical Education (NKLM) include COI (without specifically naming them) in chapter 11.1.1.2. (38). Published evidence for more comprehensive COI policies and examples for concrete COI curricula might also be suitable for the German context (39). In the US, the regular AMSA scorecard assessed COI policies at U.S. medical schools and contributed to a constant improvement in policies since its initiation in 2007 (29). Regular evaluation of the development of policies and curricula addressing COI might also be useful in Germany to incentivise and monitor progress towards better COI education at medical schools.

### 4.1 Limitations

This study is subject to several limitations. In total, 22 out of 38 medical schools did not respond to our letter and emails and therefore COI teaching activities and policies by non-responding medical schools may have been missed. Moreover, we could not validate the web-search results by Charité Universitätsmedizin Berlin, as they did not respond to extend or correct our findings. The web-search was limited to integrated search engines on the websites of medical schools and to few predefined search terms, thus the usage of a broader variety of terms and search engines may have yielded more relevant documents. Consequently, the results of this study may underestimate the number of COI policies and teaching activities that are publicly available.

Policy development is a dynamic process, some medical schools may have adopted policies after autumn 2018. When contacted, TU Dresden reported that a renewed anti-corruption directive for the medical school and university medical centre was under development. Additionally, some schools signalled willingness to introduce teaching activities and considered COI policies after we contacted them. This, however, was also the case in the study by Lieb and colleagues in 2013 (21). Our work indicates that little action was taken since then.

Medical schools don’t exist in a vacuum and further COI policies may exist at a university-wide level or at university medical centres. We argue that the consequences of COI in medicine potentially harm patient health and are therefore even more critical compared to COI that might occur in other fields. Thus, medical schools require more restrictive COI policies than other departments within a university. Teaching physicians are predominantly also employed by a university medical centre which might issue COI policies not specifically applying to medical school. However, these policies are aimed at COI of physicians working in patient care and lack specific regulations that apply to the teaching environment of medical students.

### 4.2 Conclusions

In contrast to other parts of the world, such as North America, German medical schools barely regulate students’ contact with pharmaceutical companies or teach about impacts of conflicts of interest. Several organizations (40,41) and increasingly students themselves (34–36) are demanding a cultural change in the medical profession starting with independent, unbiased medical education. COI policies at medical schools have been shown to positively impact prescribing and practise (23–25). Medical schools in Germany have a key responsibility to protect students from undue influence and enable them to critically appraise information to achieve the best possible patient care. Although national learning objectives include teaching on COI, German medical schools do too little and have a long way to go.

## Conflict of Interest Disclosure Statement

PG, LUH and DK are members of MEZIS e.V‥ SG previously consulted for Universities Allied for Essential Medicines (UAEM) Europe e.V. DK’s full disclosure is available from https://www.akdae.de/Kommission/Organisation/Mitglieder/DoI/Klemperer.pdf. The authors declare that there are no competing financial interests concerning this study.

## Acknowledgements

The authors wish to thank all participating medical schools for contributing to this study as well as the German Medical Students’ Association (bvmd e.V)., Christiane Fischer (MEZIS e.V.), Cora Koch, Paul Scheffer, Reshma Ramachandran and Barbara Mintzes for their valuable support of this project.

## Notes

http://www.interessenkonflikte.com/wp-content/uploads/2018/06/Bewertungskriterien_Interessenkonfliktstudie_Version1_42.pdf

