## Supplementary Material for "Conflict of Interest Policies at German medical schools - A long way to go"

| Medical school | Websites searched |
|---|---|
| RWTH Aachen | https://medizin.rwth-aachen.de/ |
| Universität Augsburg | https://www.med.uni-augsburg.de/<br>https://www.uni-augsburg.de/de/ |
| Charité – Universitätsmedizin Berlin | https://www.charite.de/<br>https://www.fu-berlin.de/<br>https://www.hu-berlin.de/de |
| Ruhr-Universität Bochum | http://www.medizin.ruhr-uni-bochum.de/<br>https://www.ruhr-uni-bochum.de/de |
| Rheinischen Friedrich-Wilhelms-Universität Bonn | http://ukb.uni-bonn.de/42256BC8002AF3E7/direct/home<br>https://www.uni-bonn.de/ |
| Medizinische Fakultät Carl Gustav Carus der Technischen Universität Dresden | https://tu-dresden.de/med<br>https://tu-dresden.de/ |
| Universität Duisburg-Essen | https://www.uni-due.de/med/<br>https://www.uni-due.de/ |
| Heinrich-Heine-Universität Düsseldorf | http://www.medizin.hhu.de/<br>https://www.uni-duesseldorf.de/home/startseite.html |
| Friedrich-Alexander-Universität Erlangen-Nürnberg | https://www.med.fau.de/<br>https://www.fau.de/ |
| Goethe-Universität Frankfurt | http://www.uni-frankfurt.de/54233767/fachbereich<br>http://www.uni-frankfurt.de/de?locale=de |
| Albert-Ludwigs-Universität Freiburg | http://www.med.uni-freiburg.de/de<br>http://www.uni-freiburg.de/ |
| Justus-Liebig-Universität Gießen | https://www.uni-giessen.de/fbz/fb11<br>https://www.uni-giessen.de/index.html |
| Georg-August-Universität Göttingen | https://www.med.uni-goettingen.de/index_de.php<br>http://www.uni-goettingen.de/ |
| Universitätsmedizin Greifswald | https://www.medizin.uni-greifswald.de/de/home/<br>https://www.uni-greifswald.de/ |
| Martin-Luther-Universität Halle-Wittenberg | https://www.medfak.uni-halle.de/<br>https://www.uni-halle.de/ |
| Universität Hamburg | https://www.uke.de/organisationsstruktur/medizinische-fakult%C3%A4t/index.html<br>https://www.uke.de/index.html<br>https://www.uni-hamburg.de/ |

| | |
|---|---|
| Medizinische Hochschule Hannover | https://www.mh-hannover.de/ |
| Ruprecht-Karls-Universitä t Heidelberg | http://www.medizinische-fakultaet-hd.uni-heidelberg.de/<br>https://www.uni-heidelberg.de/de |
| Universität des Saarlandes (Homburg) | http://www.uniklinikum-saarland.de/de/<br>https://www.uni-saarland.de/nc/startseite.html |
| Friedrich-Schiller-Universi tät Jena | https://www.uniklinikum-jena.de/<br>https://www.uni-jena.de/ |
| Christian-Albrechts-Unive rsität zu Kiel | http://www.medizin.uni-kiel.de/de<br>http://www.uni-kiel.de/de/ |
| Universität zu Köln | http://medfak.uni-koeln.de/<br>http://uni-koeln.de/ |
| Universität Leipzig | https://www.uniklinikum-leipzig.de/<br>https://www.uni-leipzig.de/ |
| Universität zu Lübeck | https://www.uni-luebeck.de/index.php?id=897<br>https://www.uni-luebeck.de/universitaet/universitaet.html |
| Otto-von-Guericke-Univer sität Magdeburg | http://www.uni-magdeburg.de/<br>http://www.med.uni-magdeburg.de/ |
| Johannes-Gutenberg-Uni versität Mainz | http://www.um-mainz.de<br>https://www.uni-mainz.de/ |
| Medizinische Fakultät Mannheim der Ruprecht-Karls-Universitä t Heidelberg | https://www.umm.uni-heidelberg.de/home/ |
| Philipps-Universität Marburg | https://www.uni-marburg.de/de/fb20<br>https://www.uni-marburg.de/de |
| Ludwig-Maximilians-Unive rsität München | https://www.med.uni-muenchen.de/index.html<br>https://www.uni-muenchen.de/index.html |
| Technischen Universität München | http://www.med.tum.de/de/die-fakult%C3%A4t-f%C3%B Cr-medizin/<br>https://www.tum.de/ |
| Westfälischen Wilhelms-Universität Münster | https://www.medizin.uni-muenster.de/fakultaet/start/<br>https://www.uni-muenster.de/de/ |
| Carl von Ossietzky Universität Oldenburg | https://uol.de/medizin/<br>https://uol.de/ |
| Universität Regensburg | https://www.uni-regensburg.de/medizin/fakultaet/<br>https://www.uni-regensburg.de/ |
| Universitätsmedizin Rostock | https://www.med.uni-rostock.de/<br>https://www.uni-rostock.de/ |
| Eberhard Karls Universität Tübingen | https://www.medizin.uni-tuebingen.de/de/<br>https://uni-tuebingen.de/ |

| | |
|---|---|
| Universität Ulm | https://fakultaet.medizin.uni-ulm.de/<br>https://www.uni-ulm.de/ |
| Universität Witten/Herdecke | https://www.uni-wh.de/gesundheit/department-fuer-humanmedizin/#profil<br>https://www.uni-wh.de/ |
| Julius-Maximilians-Universität Würzburg | https://www.med.uni-wuerzburg.de/startseite/<br>https://www.uni-wuerzburg.de/startseite/ |
